# Unraveling the Mechanism of a LOV Domain Optogenetic Sensor: A Glutamine Lever Induces Unfolding of the Jα Helix

**DOI:** 10.1101/2020.01.29.925040

**Authors:** James N. Iuliano, Jinnette Tolentino Collado, Agnieszka A. Gil, Pavithran T. Ravindran, Andras Lukacs, SeungYoun Shin, Helena A. Woroniecka, Katrin Adamczyk, James M. Aramini, Uthama R. Edupuganti, Christopher R. Hall, Gregory M. Greetham, Igor V. Sazanovich, Ian P. Clark, Taraneh Daryaee, Jared E. Toettcher, Jarrod B. French, Kevin H. Gardner, Carlos L. Simmerling, Stephen R. Meech, Peter J. Tonge

## Abstract

Light-activated protein domains provide a convenient, modular, and genetically encodable sensor for optogenetics and optobiology. Although these domains have now been deployed in numerous systems, the precise mechanism of photoactivation and the accompanying structural dynamics that modulate output domain activity remain to be fully elucidated. In the C-terminal light, oxygen, voltage (LOV) domain of plant phototropins (LOV2), blue light activation leads to formation of an adduct between a conserved Cys residue and the embedded FMN chromophore, rotation of a conserved Gln (Q513), and unfolding of a helix (Jα-helix) which is coupled to the output partner. In the present work, we focus on the allosteric pathways leading to Jα helix unfolding in *Avena sativa* LOV2 (AsLOV2) using an interdisciplinary approach involving molecular dynamics simulations extending to 7 μs, time-resolved infrared spectroscopy, solution NMR spectroscopy, and in-cell optogenetic experiments. In the dark state, the side chain of N414 is hydrogen bonded to the backbone N-H of Q513. The simulations predict a lever-like motion of Q513 after Cys adduct formation resulting in loss of the interaction between the side chain of N414 and the backbone C=O of Q513, and formation of a transient hydrogen bond between the Q513 and N414 side chains. The central role of N414 in signal transduction was evaluated by site-directed mutagenesis supporting a direct link between Jα helix unfolding dynamics and the cellular function of the Zdk2-AsLOV2 optogenetic construct. Through this multifaceted approach, we show that Q513 and N414 are critical mediators of protein structural dynamics, linking the ultrafast (sub-ps) excitation of the FMN chromophore to the microsecond conformational changes that result in photoreceptor activation and biological function.

## Introduction

The C-terminal light, oxygen, voltage (LOV) domain from plant phototropins are versatile protein domains that have been adapted for protein engineering and molecular imaging.^1,2^ LOV photoreceptors are members of the Per-ARNT-Sim (PAS) domain superfamily and use a non-covalently bound flavin mononucleotide (FMN) cofactor to sense 450 nm (blue) light (**Figure 1A**).^3^ Light excitation initiates a photocycle in which a singlet excited state undergoes intersystem crossing to a triplet excited state which subsequently forms a covalent Cys-FMN-C4a adduct on the μs timescale that absorbs at 390 nm (A390).^4^ Formation of the A390 Cys adduct and accompanying protonation of the adjacent N5 position^5^ are thought to be the driving force behind the structural changes that accompany effector activation in the LOV domain family including alterations in the structure and conformation of the C-terminal Jα helix.^6^

**Figure 1:**
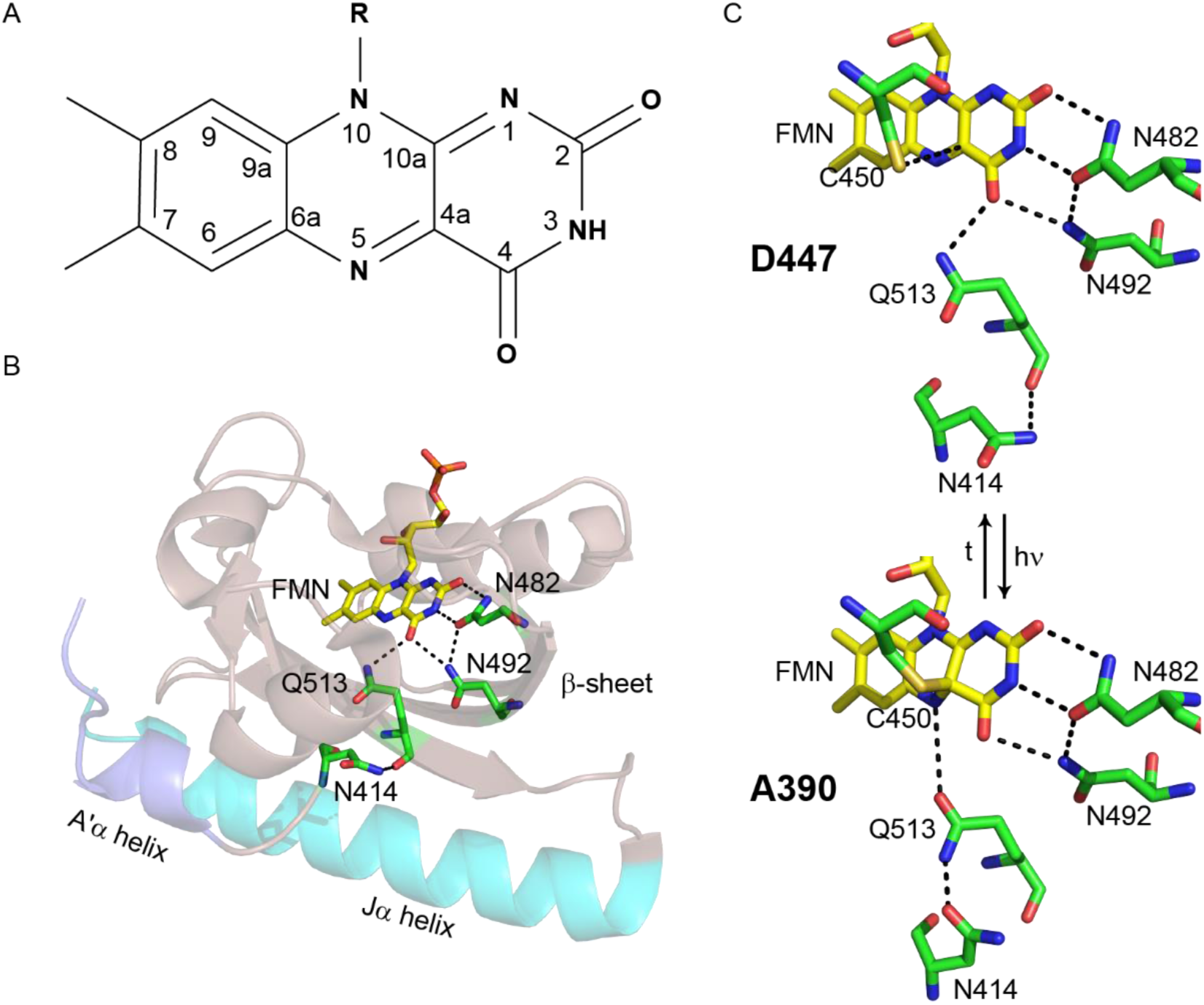
The AsLOV2 domain. (A) The isoalloxazine ring of the FMN cofactor. (B) The FMN cofactor (yellow) makes hydrogen bonding interactions with Q513, N492 and N482. Q513 in turn is hydrogen bonded to N414. The Jα helix is shown in cyan and the A’α helix is shown in slate. (C) The hydrogen bonding network around FMN is shown for both the dark (D447) and light (A390, adduct) states. The flipped conformation of Q513 is shown for the light state structure. The figure was made using PyMOL Molecular Graphics Software^7^ using the crystal structures of AsLOV2 (PDB 2V1A (dark), 2V1B (light)).^8^

The LOV2 domain from *Avena sativa* phot1 (AsLOV2) is a model system for LOV photoreceptor activation (**Figure 1B**). The isoalloxazine ring of the FMN cofactor is surrounded by a hydrogen-bonding network that senses and responds to excitation on the ultrafast timescale leading to activation of a Ser/Thr kinase that regulates phototropism in plants.^9^ Photoexcitation results in formation of an adduct between C450 and FMN and protonation of FMN-N5 which results in rotation of a conserved Gln (Q513) and unfolding of a conserved C-terminal α helix (Jα helix).^9^ Structural changes are also observed in the LOV domain β-sheet,^10,11^ and an N-terminal helix (A’α) is thought to act as a regulatory element which unfolds concurrently with changes in the Jα helix.^12,13^

Optogenetics and optobiology are emerging fields in which a range of biological functions can now be controlled by light. The LOV domain was first used in an optogenetic construct by Möglich and coworkers, in which the LOV domain of YtvA was fused to a histidine kinase (YF1).^14^ Fusion proteins utilizing AsLOV2 followed, with the development of the photo-activatable LOV-Rac1 sensor (PA-Rac1) in which the Jα helix was used as a reversible photocage controlling the activity of the GTPase Rac1.^15,16^ Further protein engineering led to the development of iLID, which added light inducible dimerization capabilities and a recognition peptide embedded within the sequence of the Jα helix such that unfolding of the Jα helix enabled recruitment of signaling proteins in-cell,^17^ and LEXY, which included a nuclear export sequence in the Jα helix so that nuclear export could be controlled by blue light.^18^ Despite these advances, some LOV-based optogenetic tools still possess a significant level of activity in the dark state, resulting in sub-optimal dynamic range and limiting their broad deployment. A complete molecular understanding of the mechanism of LOV domain function is thus required for the rational optimization of LOV-based optogenetic photoreceptors.^19,20^

Molecular dynamics (MD) simulations have previously guided hypotheses into the mechanism of formation,^21^ and breakdown,^22^ of the Cys-FMN adduct, and the accompanying structural dynamics,^13,23^ in native LOV domains and mutants lacking the Cys residue.^24^ Initial studies by Peter et al. focused on the role of the Iβ and Hβ strands in signal propagation,^23^ resulting in the proposal that stress on Iβ leads to rearrangement of the hydrogen bonding contacts between Hβ and the Jα helix and helix unfolding. It was also proposed that the FMN binding pocket undergoes dramatic changes in which N482 and N492 move out of the pocket to maintain contacts between Q513 and FMN. Freddolino et al. extended this work by increasing the simulation timescale to 1 µs,^13^ leading to the identification of a potential salt bridge between the A’α and Jα helices, and supporting previous studies of an interaction between the two helices. The simulations also suggested that the N414 side chain N-H forms a H-bond with the Q513 side chain C=O during light state formation. The role of N414 in photoactivation has been tested experimentally in which the N414A, N414V and N414Q mutations modulate the cycling time of AsLOV2 with time constants of 1427 s, >720 s, and 280 s, respectively compared to 80 s for wild-type.^25^ In the case of N414V, a slightly unfolded Jα helix was observed in the dark state.^12^

LOV structural dynamics have been studied extensively using infrared spectroscopy and more recently time-resolved infrared spectroscopy (TRIR). A marker band for unfolding of the Jα helix was identified in the amide I region, ∼1620-1640 cm^−1^ using difference FTIR.^26–28^ TRIR experiments revealed that helix unfolding was multi-step,^29^ and that structural changes resulting from adduct formation follow dispersive kinetics commonly associated with sampling of multiple protein conformations prior to reaching a metastable state, which is the signaling state in LOV domain proteins.^30^ TRIR spectroscopy of multiple LOV domain proteins revealed variations in the dynamics of the β-sheet and the Jα helix, which link the LOV domain to the relevant effector domains.^31^ Despite the information obtained from time-resolved spectroscopy, the mechanisms of signal propagation from FMN to the effector domain are still largely unknown.

Using a combination of theoretical and experimental approaches, we have now directly probed the evolution of the structural dynamics in AsLOV2 that couple flavin photoexcitation with structural changes at sites that are remote from the chromophore. In particular, we have determined that the rotation of Q513 out of the flavin binding pocket and the formation of a transient hydrogen bond with N414 is a key step in the mechanism of Jα helix unfolding. Site directed mutagenesis combined with time-resolved infrared spectroscopy and NMR spectroscopy was used to interrogate this mechanism and we found that the Jα helix was partially unfolded in the dark state and unfolding kinetics were accelerated in the N414A mutant and delayed in N414Q AsLOV2. To correlate photoreceptor dynamics and function, the impact of the mutants was assessed at the cellular level in the Zdk2-AsLOV2 dimerization (LOVTRAP) system where it was found that the N414A mutant showed a 4-fold reduction in activity and dynamic range when exposed to blue light.

## Materials and Methods

### Molecular Dynamics Simulations

Molecular dynamics (MD) simulations were used to analyze the structural dynamics of AsLOV2 starting from the X-ray structure of AsLOV2 in the light state. Solution NMR spectroscopy has shown that the AsLOV2 Jα helix is unfolded in the light state. However, this helix is still folded in the X-ray structure of the AsLov2 light state which is thought to be due to crystal contacts between protein molecules. Thus, the dark state X-ray structure is an ideal starting point for monitoring Jα helix unfolding since the unfolded helix should be the most stable species in solution.

### Parameter Development for the Flavin Cofactor

Parameter generation proceeded in 2 parts. First, a library file corresponding to the flavin alone was created. Coordinates were extracted from protein data bank (PDB) 2V1B for AsLOV2 in the light state.^1^ All atoms were deleted except for those of the flavin. Chimera was used to add hydrogen atoms, and the hydrogen atom added to atom C4A was then deleted since this is the site of Cys adduct formation.^2^ These coordinates were loaded into the Amber antechamber module using a charge of −3 to generate AM1BCC partial charges and assign atom types for use with the GAFF2 general force field.^3^ The resulting mol2 file was loaded into Amber parmchk2 to create a frcmod file, which was then loaded with the mol2 file into Amber tleap to create an Amber library file for the flavin. This provided the library files appropriate for the structure of the flavin residue, ready to connect to the protein at C450. However, the force field parameters do not correspond to the state following formation of the adduct after light activation and additional steps were needed.

The next step involved generation of force field parameters for the flavin in the light-activated adduct state. The process above was repeated except that the coordinates retained from 2V1B included the flavin as before, along with the SG and CB atoms from C450. Chimera was used to add hydrogen atoms, resulting in a methyl capping group at the position of the C450 CB. Antechamber was used to generate AM1BCC partial charges and GAFF2 atom types. Parmchk2 and tleap were used to create the Amber library file for the flavin attached to the SG and CB. Then, the partial charges and atom types obtained in this step were copied to the atoms in the library file obtained using only the flavin (described above), such that the final library file included only flavin atoms, ready to accept a bond from C450, but with atom types and partial charges appropriate for the state following adduct formation with C450. The bond angle and dihedral parameters used were those obtained from GAFF2 using the flavin connected to the SG and CB atoms. Force field parameters for the GAFF2↔ff14SB interface across C4A-SG were obtained by adopting the parameters assigned by GAFF2 using the flavin+SG+CB fragment. This parameter file was edited to change the atom types for the SG and CB atoms from those obtained using antechamber to those appropriate for Cys in the ff14SB protein force field (c3 → 2C, ss → S). In this manner the protein force field was applied inside Cys, but parameters for the Cys-flavin linkage (FMN-C) were adopted from GAFF2. Finally, since the Cys backbone CA atom was not present in the larger flavin fragment but is connected to the flavin through a dihedral term, parameters for the dihedral C3-S-2C-CX (corresponding to atom names C4A-SG-CB-CA) were adapted from C3-S-C3-C3 in GAFF2. The resulting library file and force field parameter file for the flavin in the light-activated state are included as Supporting Information.

### Initial Modeling of FMN-C into the Dark State Structure

Coordinates for AsLOV2 in the dark state were obtained from PDB ID 2V1A.^1^ Glycerol molecules were removed and 143 structured water molecules were retained. Hydrogen atoms were added and the system was solvated in a truncated octahedral box using a buffer of 8 Å minimum between any solute atom and the box boundary, resulting in addition of 5014 water molecules. The OPC 4-point water model was employed,^4^ the parameters described above were used for the flavin, and the ff14SBonlysc protein force field was used.^5^ This model includes the side chain updates from ff14SB,^5^ but not the empirical backbone corrections included in that model for use with TIP3Pwater.^6^ A covalent bond was added between the SG atom of C450 and the C4A atom of the flavin, corresponding to the bond that forms upon light activation.

### Equilibration of the FMN-C AsLOV2 Model

Minimization and equilibration were carried out using Amber version 16.^7,8^ Initial minimization was performed for 100000 steps with restraints on all atoms except hydrogens, water, C450 and the flavin. The restraint force constant was 100 kcal·mol^−1^·Å^−2^. Next, the system was heated from 100 K to 298 K over 1 ns in the NVT ensemble, with a time step of 1 fs and SHAKE on all bonds including hydrogen. A nonbonded cutoff of 8 Å was used, with long-range electrostatics included by the particle mesh Ewald method.^9^ The same restraints were maintained. Temperature was maintained using a Langevin thermostat with γ set to 1.0. Next, pressure and density were relaxed using 1ns NPT simulation at 298 K with a strong pressure coupling constant of 0.1 and all other parameters maintained from the prior step. Next, 1 ns MD was performed using the same protocol but with restraint force constant reduced to 10.0 and pressure coupling constant increased to 0.5. Next, minimization was performed for 10000 steps after removing all restraints except those on protein backbone CA, N and C atoms and no restraints on C450. Next, 1 ns MD in the NPT ensemble at 298 K was performed using the same protocol as above, with restraints only on backbone CA, N and C atoms excepting C450. The restraint force constant was then reduced from 10.0 to 1.0 and an additional 1 ns MD was carried out, followed by 1 ns MD with restraint force constant reduced to 0.1. Finally, 1 ns fully unrestrained MD was performed with NPT at 298 K.

### Production runs

Production runs followed the same protocols as equilibration, except that hydrogen mass repartitioning was used to enable a 4 fs time step as has been described elsewhere.^10^ The simulation was performed on NVIDIA GPUs using the CUDA version of Amber.^11^ MD was continued until approximately 7.5 microseconds of dynamics were generated.

### Analysis

Analysis of the MD simulation output was performing using the cpptraj module of Amber,^12^ along with custom python scripts.

### Site-directed Mutagenesis

The N414A and N414Q mutations in AsLOV2 were created in the pET15b-AsLOV2 and pHis-Gβ1-Parallel-AsLOV2 constructs using the QuickChange method and KOD HotStart polymerase (Novagen).

### Expression and Purification of AsLOV2 for TRIR

AsLOV2 and N414 mutants were expressed and purified as described previously.^13^ BL21(DE3) competent cells were transformed by heat shock at 42°C with pET15b-AsLOV2 (wild-type or mutant) and the plated on LB-Agar plates containing 200 µg/mL ampicillin. Following overnight incubation at 37°C, single bacterial colonies were used to inoculate 10 mL of 2X-YT media containing 200 µg/mL ampicillin. After overnight incubation at 37°C in an orbital shaker (250 RPM), the 10 mL cultures were used to inoculate 1 L of 2X-YT media supplemented with 200 µg/mL ampicillin. The cultures were incubated at 37°C in an orbital shaker (250 RPM) until the OD_600_ reached ∼0.8-1.0 at 37°C and protein expression was then induced by addition of 1 mM IPTG (Gold Biosciences). Protein expression was allowed to proceed overnight at 18°C in an orbital shaker (250 RPM).

Cells from the 1 L cultures were harvested by centrifugation at 5,000 RPM (4°C) and resuspended in 40 mL of 20 mM Tris buffer pH 8.0 containing 150 mM NaCl. The resuspended cells were lysed by sonication and cell debris was removed by ultracentrifugation at 40,000 RPM for 1 h (4°C). The clarified lysate was the loaded onto a 5 mL HisTrap FF column equilibrated with resuspension buffer. The column was then washed with 100 ml of resuspension buffer containing 20 mM imidazole, and protein was eluted with resuspension buffer containing 500 mM imidazole. Fractions containing protein were pooled, concentrated to a volume of 5 mL using a 10 kDa MWCO concentrator (Amicon), and loaded onto a Superdex-75 column equilibrated with 20 mM Tris buffer pH 7.6, containing 150 mM NaCl. The protein was concentrated to 200 µM using 10 kDa MWCO concentrator and lyophilized for storage and prior to exchange into D_2_O for the TRIR measurements. Purity was >95% by SDS-PAGE.

### Expression and purification of 15N labelled AsLOV2

Isotope labeling of AsLOV2 was performed using the pHis-Gβ1-Parallel1 expression vector containing the DNA encoding wild-type AsLOV2 or the N414A or N414Q mutants (pHis-Gβ1-Parallel-AsLOV2). Protein expression and purification were performed as described previously with some modifications.^14^ Briefly, BL21(DE3) cells were transformed by heat shock at 42°C with pHis-Gβ1-Parallel-AsLOV2 and plated on LB-agar plates supplemented with 100 µg/mL ampicillin (Gold Biosciences). After incubation overnight at 37°C, a single colony was used to inoculate 10 mL of 2X-YT media containing 100 µg/mL ampicillin. After overnight incubation at 37°C in an orbital shaker (250RPM), cells were harvested by centrifugation and rinsed by resuspension in 2x 10 mL volumes of M9 salts followed by centrifugation. The rinsed cells were then used to inoculate 1 L of M9 medium supplemented with 1 g/L ^15^N-NH_4_Cl (Cambridge Isotope Labs), 4 g/L dextrose, 1X MEM vitamins, 1 mM MgSO_4_, 10% glycerol, and 100 µg/mL ampicillin. Bacterial cultures were incubated at 37°C in an orbital shaker (250 RPM) until the culture reached an OD_600_ of ∼0.6-0.8. Protein expression was induced by adding IPTG to a final concentration of 1 mM followed by incubation at 18°C overnight (14-18 hours) in an orbital shaker (250 RPM).

Bacterial cells were harvested by centrifugation at 5,000 RPM for 10 min at 4°C, and the cell pellet was subsequently resuspended in 20 mM Tris buffer pH 8.0 containing 150 mM NaCl buffer. Cells were lysed by sonication and cell debris was removed by ultracentrifugation at 40,000 RPM for 1 h and 4°C. All subsequent purification steps were carried out on an AKTA FPLC (Pharmacia Biosciences). The clarified lysate was loaded onto a 5 mL HisTrap FF Ni-NTA column (GE) equilibrated with the resuspension buffer. The column was subsequently washed with resuspension 100 mL buffer containing 20 mM imidazole, and protein was eluted using resuspension buffer containing 500 mM imidazole. Fractions containing protein were pooled and desalted to remove imidazole by gel filtration on a 50 mL BioScale P-6 column (BioRad) using resuspension buffer as the eluant.

The Gβ1-His tag was removed using 1 mg of His-TEV protease per 30 mg of protein. Gβ1-His tag AsLov2 was incubated with His-TEV protease overnight at 4°C on a rocking platform. The Gβ1-His tag and His-TEV were separated from AsLOV2 using a 5 mL HisTrap FF column equilibrated with 20 mM Tris buffer pH 8.0 containing 150 mM NaCl. Fractions containing AsLOV2 were collected and pooled to give a volume of XX mL, and the protein was then exchanged into 50 mM sodium phosphate buffer pH 7.0 containing 150 mM NaCl and 6 mM sodium azide by size-exclusion chromatography on Superdex-75 16/60 column equilibrated with the phosphate buffer. The protein was concentrated to 500 µM using a 10 kDa MWCO concentrator for the NMR spectroscopy experiments. Protein purity was >95% by SDS-PAGE.

### Time-Resolved Multiple Probe Spectroscopy (TRMPS)

TRIR spectroscopy was carried out at the Central Laser Facility (Harwell, UK) using the Time resolved multiple probe spectroscopy approach (TRMPS) which has been described previously.^15,16^ Mid-IR probe light was generated using an OPA with a DFG stage pumped by a Ti:Sapphire laser and the signal and idler outputs were mixed to generate the broadband probe centered at ∼1550 cm^−1^ with a pulse duration of <100 fs at a repetition rate of 10 kHz. The 450 nm visible pump was generated by a second Ti:Sapphire laser pumped OPA (SpectraPhysics Ascend and Spitfire) with a pulse duration of ∼100 fs and a 1 kHz repetition rate. Probe light was detected using two 128-pixel MCT detectors to give ∼400 cm^−1^ spectral bandwidth. The spectra were calibrated to polystyrene film giving a resolution of 3 cm^−1^. The approach used in this work was modified to include a chopper to modulate the repetition rate of the pump laser from 1 kHz to 500 Hz such that a pump-off subtraction of the baseline was performed on each spectrum prior to excitation, eliminating fixed pattern noise and enabling longer time delay acquisitions.^17^ Protein samples were concentrated to 1 mM using a 10 kDa MWCO concentrator and ∼1 mL of this solution was loaded into a Harrick cell modified for low volume flow at a rate of 1 mL/min. A 50 µm spacer was used between two CaF_2_ windows and the sample cell was rastered to ensure fresh sample for each laser shot.

### ^1^H-^15^N HSQC NMR

Multidimensional ^1^H-^15^N HSQC NMR spectroscopy was performed on a Bruker Avance III HD 800 MHz Spectrometer. Both dark and light state spectra of wild-type and mutant AsLOV2 proteins were acquired as previously described.^14^ The light state of AsLOV2 was generated by illumination using a Coherent Sapphire Laser (488 nm, ∼200 mW) focused into a fiber optic wand that was submerged into the protein solution in the NMR tube. The power at the end of the fiber was set to 50 mW and a shutter controlled by Bruker Topspin Software enabled pulses of 120 ms prior to each transient.^18^ Processing and analysis was performed using Bruker TopSpin Software.

### LOVTRAP Cellular Assay

InFusion cloning (Clontech) was used to attach iRFP to the N-terminus of AsLOV2 (wild-type, N414A or N414Q mutants). The pHR lentivirus vector, ZDK insert and AsLOV2 insert sequences were amplified using HiFi or GXL polymerase, subjected to infusion cloning, and amplified using Stellar competent cells. Sequences were verified using restriction digests and Sanger sequencing (GeneWiz).

Lenti-X 293T cells were used to generate virus for transfection and expression of constructs. Cells were maintained in a 6-well plate containing 2 mL of DMEM supplemented with 10% FBS, 100 µg/mL penicillin/streptomycin, and 2 mM L-glutamine per well or T25 flasks containing 5 mL of the same medium at 37° with 5% CO_2_.

Lentiviral particles containing the iRFP-AsLOV2 construct or the ZDK p2a Fusion Red construct were generated by transfecting Lenti-X 293T cells with the pHR-iRFP-AsLOV2 vector or the pHR-ZDK-p2a-FusionRed together with the pCMV-dR8.91 and pMD2.G lenti-helper plasmids in a 1.5:1.3:0.17 ratio, respectively. The helper plasmids were gifts from the Torono lab via Addgene, and transfections were performed using Fugene transfection reagent (Promega). The transfected Lenti-X 293T cells were incubated at 37° with 5% CO_2_ for two days, after which the media containing lentivirus particles was filtered through a 0.45 µm filter. Forty µL of 1 M HEPES buffer pH 7.4 was then added to ∼2 mL of the filtered media which was then stored at 4°C for two weeks for immediate use or at −80°C for long term storage.

Prior to infection, LentiX 293Ts were plated at 50% confluency and incubated overnight to attach. Five hundred µL of the viral media was then added to the cells. After incubation for 24 h at 37°C with 5% CO_2_, the cells were washed with fresh media and then incubated at 37°C with 5% CO_2_ for another 24 h. Cells to be imaged were plated 12 h prior to imaging in 96-well, black-walled, glass bottomed plates (InVitro Scientific) coated with 100 µl of 10 µg/mL fibronectin diluted in PBS which was rinsed off with PBS prior to plating cells in full media. Confocal microscopy was performed on a Nikon Eclipse Ti Microscope equipped with a linear motorized stage (Pior), CSU-X1 spinning disk (Yokogawa), using 561 nm (ZDK-p2a-FusionRed) and 650 nm (iRFP-AsLOV2) modules (Agilent) laser lines. Images were acquired using a 60x oil-immersion objective and an iXon DU897 EMCCD camera. For FMN excitation, 447 nm light was provided by a LED light source (Xcite XLED1) through a digital micromirror device (Polygon400, Mightex Systems) to temporally control light inputs.

## Results

The objective of this work was to characterize the structural dynamics that lead to Jα helix unfolding in AsLOV2. To accomplish this goal we used MD simulations to analyze the structural changes that accompany light state formation by inserting the Cys-FMN light state adduct into the dark state structure of AsLOV2 and then allowing the MD simulations to ‘relax’ the protein structure to adjust to the presence of the Cys-FMN adduct. The MD simulations were used to identify key residues implicated in the pathway leading to Jα helix unfolding. Site-directed mutagenesis was then used to probe the role of these residues in light state formation and AsLOV2 dynamics using a combination of MD simulations, static and time-resolved infrared spectroscopy, ^15^N/^1^H-HSQC NMR spectroscopy, and cell-based optogenetic experiments. Based on this analysis we propose a mechanism for photoactivation in which a hydrogen bonding interaction between the NH side chain of N414 and backbone C=O of Q513 is broken and a transient hydrogen bond is formed between the NH and C=O side chain groups of N414 and Q513, respectively. Q513 is thus shown to act as a lever which forces the Jα helix away from the LOV domain thereby initiating helix unfolding and activation of the downstream effector module.

### Molecular dynamics (MD) simulations of AsLOV2

To identify allosteric pathways leading to unfolding of the Jα helix in AsLOV2, we used MD simulations to predict structural changes in response to Cys-FMN formation. The Cys-FMN adduct was taken from the light state crystal structure (PDB 2V1B),^8^ and parameterized prior to insertion into the dark state crystal structure (PDB 2V1A). The dark state crystal structure containing the light state Cys-FMN adduct was equilibrated prior to the MD simulations which were then performed in 4 fs steps for >7 μs to simulate the response of the protein to Cys-FMN adduct formation.

The data from the simulations are presented as a heat map in **Figure 2A** with time on the x-axis, residue number on the y-axis, and a color bar to show increasing root mean squared deviation (RMSD, log scale) from the initial dark state structure. The simulation reveals that residues in a loop immediately preceding the Jα helix become perturbed by adduct formation early in the simulation and propagate through the N-terminal end of the Jα helix with the most dramatic changes in the helix occurring at ∼5 μs with complete loss of helical character.

**Figure 2:**
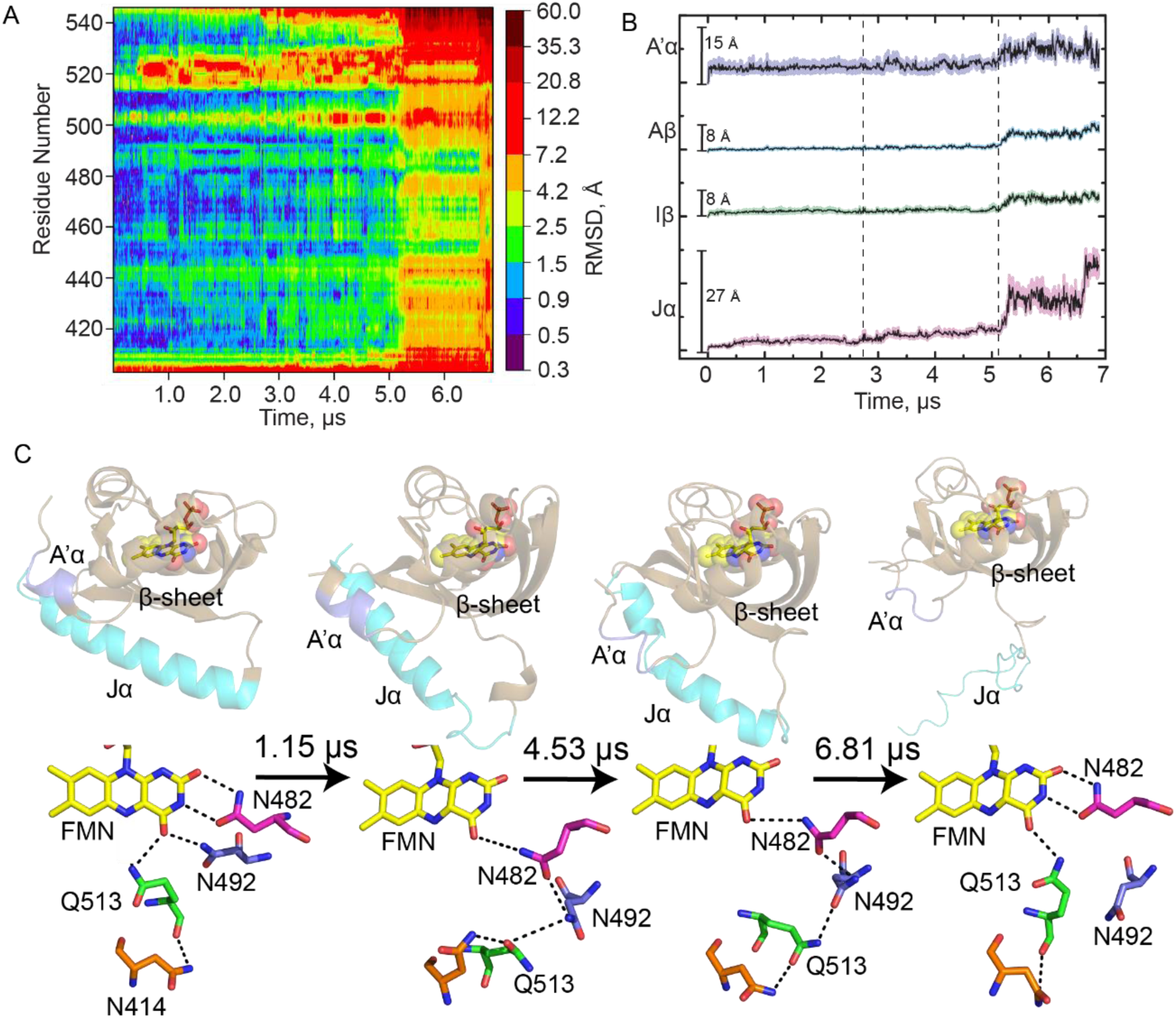
Molecular dynamics simulations reveal hydrogen bonding pathways leading to Jα helix unfolding. (A) RMS as a function of time post-adduct formation for each residue in AsLOV2 is shown as a heat map with increasing simulation time on the x-axis and residue number on the y-axis. A color bar shows increasing RMS in Å from the initial dark state structure. (B) Average RMS ± SEM for select structural components of the LOV domain: A’α helix (slate), Aβ (cyan), Iβ (green), and Jα (pink). (C) The evolution in secondary structure is shown together with changes in hydrogen bonding interactions between the flavin and residues N482, N492, Q513 and N414 for simulation times of 0 (dark state AsLOV2, 2V1A), 1.15 μs, 4.53 μs, and 6.81 μs (Light state, MD) after adduct formation. Hydrogen bonds are shown as black dotted lines. The figure was made using PyMOL Molecular Graphics Software^7^ using predicted structures from MD simulations performed in Amber.^8,32^

Averaged RMS traces as a function of time are shown for several key regions of AsLOV2 with maximum RMS shown as a scale bar (black line, **Figure 2B**). The standard error of the mean (SEM) is shown as shaded color around the black line. The A’α helix (residues 404-410, slate) shows the most variability very early in the simulation with both high average RMS and SEM, consistent with previous experimental work.^12^ The RMS increases again at 5 μs, on the timescale of unfolding of the Jα helix. RMS for the β-sheet (Aβ (residues 414-418, cyan) and Iβ (residues 506-516, green)) are also shown and predict significant perturbations after 5 µs. The Jα-helix (residues 522-544, pink) shows the most changes in RMS over time, as expected for the helical to disordered transition that has been shown to occur upon light state formation by Gardner and others.^9,33^ The unfolding of the Jα helix appears to occur in four phases with increases in RMS at 0.5 μs, 2.75 μs, 5.25 μs, and 6.60 μs. The 0.5 μs phase involves the initial undocking of the Jα-helix from the β-sheet (**Figure 2C, S1**) and the formation of a helical structure in the loop adjacent to the Jα helix (1.15 μs snapshot). The 5.25 μs phase is assigned to more significant disordering of the Jα helix which results in complete unfolding of the helix by ∼6.60 μs.

The changes in secondary structure are accompanied by specific changes in the residues that surround the FMN chromophore. In the dark state N414 is hydrogen bonded to the backbone amide of Q513. By the 0.38 μs time point of the simulation Q513 is predicted to move 2.4 Å and 5.7 Å from its initial backbone and side chain positions, respectively, and rotate 62° out of the binding pocket. This motion of Q513 pulls the β-sheet such that the side-chains of N482 and N492 lose their hydrogen bonding interactions with C2=O, C4=O, and N3-H of FMN. In the 4.53 µs structure, N482 appears to undergo a 3.1 Å slide out of the pocket while N492 rotates outward by 31° resulting in a side chain motion of 3.7 Å. These residues appear to form a chain that includes N414, Q513, N492, and N482 in which N482 remains hydrogen bonded to the flavin C4=O (Figure 2C, 1.15 μs and 4.53 μs structures), consistent with simulations by Freddolino et al.^13^ N482 appears to move back toward C2=O over time while N492 undergoes a rotation along the protein backbone and remains rotated out and exposed to solvent (between 1.15 μs to 6.81 μs). In the 6.81 μs structure, N482 has returned to its initial dark state position while N492 is predicted to remain rotated out of the pocket, 4.4 Å from its initial position. Q513 is shown to be hydrogen bonded to C4=O, though in a slightly different orientation compared to that observed in the X-ray structure of the light state (**Fig S2**). The differences can be explained by the method of generating light state crystals in which the protein crystals of the dark state were illuminated prior to data collection and may not reflect the flexibility of the protein observed in solution measurements. Simultaneously, N414 is pulled away from Q513 due to its proximity to the A’α helix which becomes disordered prior to unfolding of the Jα helix. We propose that the transient hydrogen bond between N414 and Q513 links A’α to FMN C4=O via a 15 Å signal transduction wire also involving N482 and N492 thereby inducing unfolding of the A’α helix concomitant with distortion of the β-sheet such that the Jα helix is pushed far enough away from the LOV core to induce its unfolding.

### A transient hydrogen bond between Q513 and N414 modulates helix unfolding

The MD results were used to inform fs-ms TRIR measurements, suggesting transient studies of N414A and N414Q-AsLOV2 mutants could provide valuable insight into Jα helix unfolding dynamics. The N414A mutation was chosen to remove potential hydrogen bond interactions with the N414 side chain and the N414Q mutation was chosen to retain an amide side chain potentially capable of forming a hydrogen bond with Q513.

We first measured the TRIR spectra of wild-type AsLOV2 using time resolved multiple probe spectroscopy (TRMPS, **Figure 3A**). The band assignments and time constants of 2.3 ns, 8.8 μs, and 14 μs corresponding to the excited state decay, triplet state decay, and A390 formation are consistent with the global analysis of our previous data from Gil et. al. (**Figure 3B**).^30^ A fourth, 313 μs time constant (EAS5) has been added to the global fitting model for wild-type AsLOV2 which describes the formation of the final signaling state as observed in light minus dark (L-D) steady state FTIR measurements (**Figure 3B; Figure S4** shows the comparison between EAS5 and L-D FTIR). This additional EAS5 is termed “Sig” and corresponds to full disordering of the helix and light state formation. The bleach at 1626 cm^−1^, which corresponds to disorder of the Jα helix, reaches a maximum of −0.35 mOD on this timescale and does not recover within the time resolution of the experiment (**Figure 4A**). The transient at 1636 cm^−1^ shows a rise and decay on the µs timescale assigned to an intermediate state between adduct formation and Jα helix unfolding, and consistent with the presence of a transient hydrogen bond from the MD simulations. Therefore, the full activation pathway of AsLOV2 can be probed using the TRMPS method.

**Figure 3:**
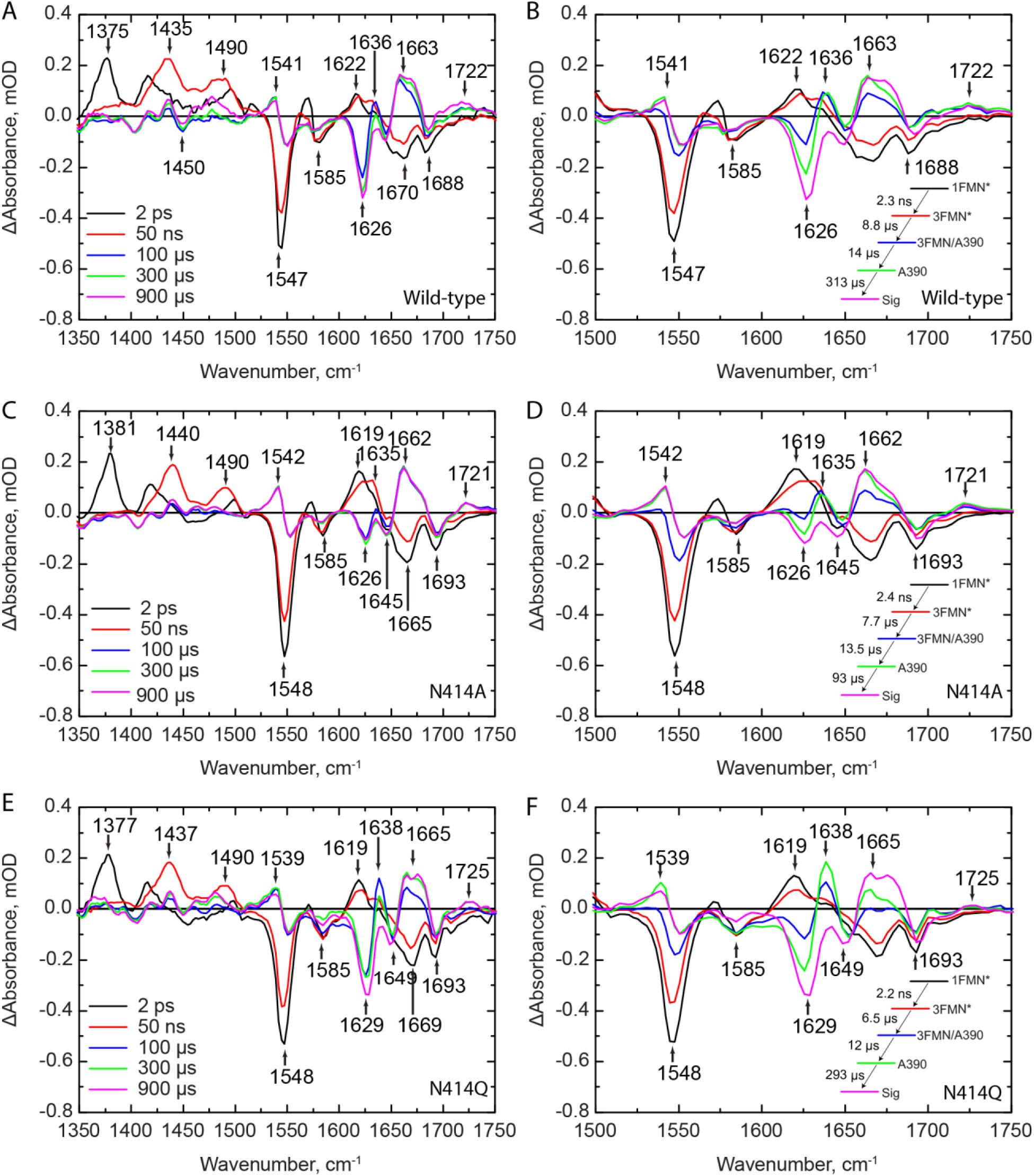
TRIR spectra of wild-type and mutant AsLOV2 proteins shows reduced helix unfolding in N414A-AsLOV2. TRIR spectra for (A) wild-type, (C) N414A, and (E) N414Q-AsLOV2 are shown at time delays of 2 ps, 50 ns, 100 μs, 300 μs, and 900 μs. The EAS from a global fit to a sequential exponential model are shown adjacent to the respective TRIR panels B, D, and F, respectively. While the excited and triplet state decays are not affected by mutation of N414, there is a 3-fold acceleration in the rate of formation of the final Sig state in N414A AsLOV2. The N414 -AsLOV2 mutant also shows reduced changes in the 1626 cm^−1^ band assigned to the Jα helix while N414Q AsLOV2 shows a reduced rate of structural dynamics and larger amplitude of the 1638 cm^−1^ band in the A390 state.

**Figure 4:**
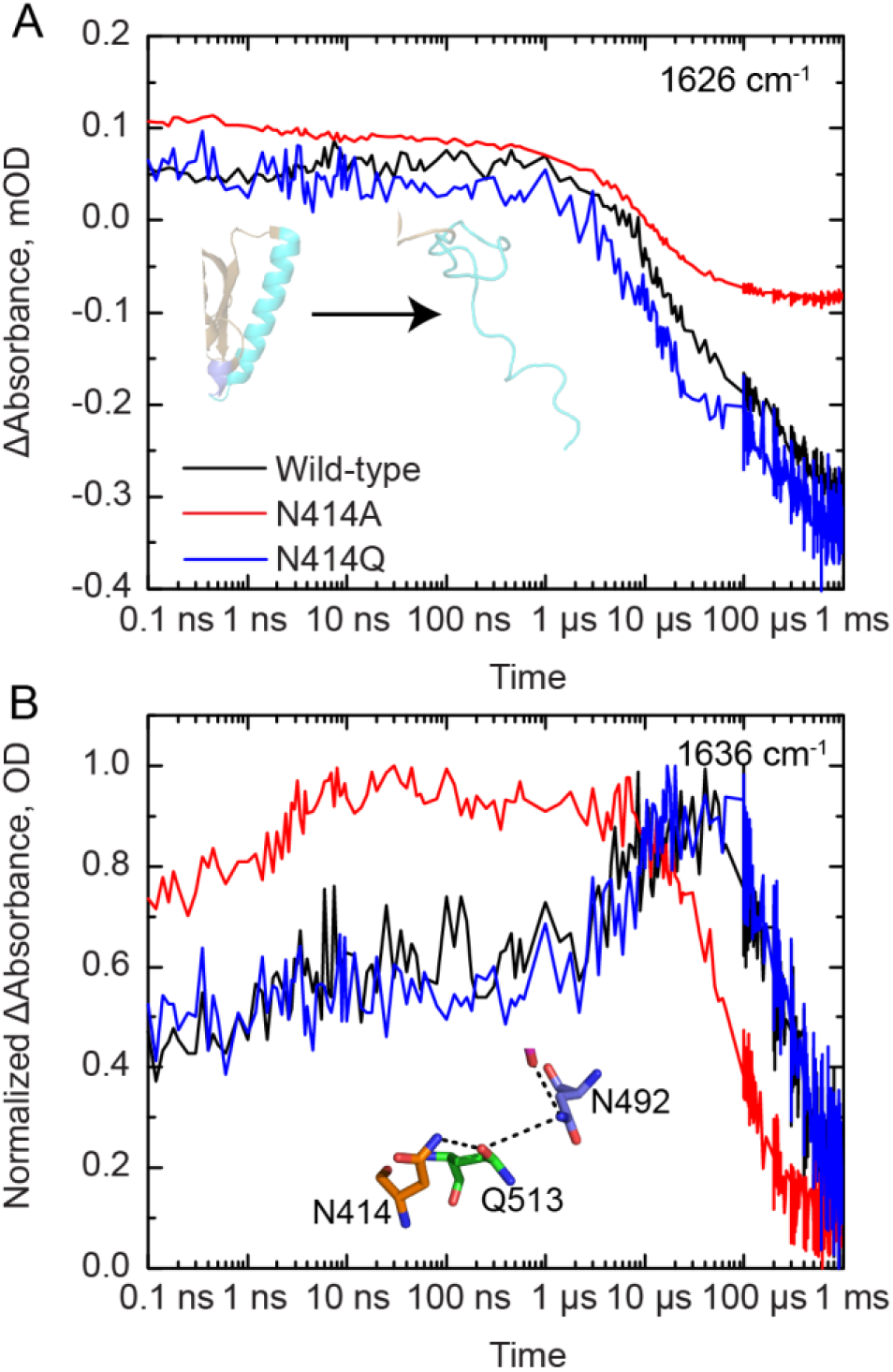
Selected kinetic traces from TRIR spectra of wild-type and mutant AsLOV2. (A) The unfolding of the Jα helix is tracked by the increase in bleach intensity at 1626 cm^−1^ for the wild-type (black), N414A (red) and N414Q (blue) AsLOV2 proteins. (B) A rise and decay of the signal at 1636 cm^−1^ is assigned to structural dynamics associated with a transient hydrogen bond between N414 and Q513 due to the rotation of Q513. This signal decays to zero with the time constant of the fourth EAS.

We then measured the TRIR spectra of N414A and N414Q AsLOV2 (**Figure 3C,E**). Whilst the excited state (^1^FMN* and ^3^FMN*) spectra and kinetics are identical to those of the wild-type protein, the changes to N414 result in dramatic differences in the A390 to Sig EAS. In N414A AsLOV2 (**Figure 3C,D**), the bleach at 1626 cm^−1^ corresponding to the disordered Jα helix is reduced ∼3-fold in intensity to −0.12 mOD, suggesting that the changes in Jα are reduced and partially decoupled from chromophore activation (**Figure 4A, red trace**). The transient at 1635 cm^−1^ is still evolving in a similar manner to that observed in the wild-type protein, although the rise in the transient is not as pronounced in the N414A mutant **(Figure 4B**). In addition, the time constant for evolution of the A390 to Sig states is accelerated ∼3-fold to 93 μs compared to the 313 μs time constant determined for wild-type AsLOV2. This suggests that the final “Sig” state forms faster in N414A AsLOV2, further suggesting altered structural dynamics in this mutant.

In the N414Q mutant, the amide side chain of N414 is conserved by mutation to Q414, preserving the potential for formation of a transient hydrogen bond. The EAS are shown in Figure 3F. While overall very similar to the wild-type AsLOV2, there are several results that deserve comment. The time constant determined for the A390 to Sig EAS is 293 μs, which is very similar to that of the wild-type at 313 μs. However, the transient at 1636 cm^−1^ in N414Q is larger than that of the wild-type as shown in the raw μs-timescale data (Figure 3F), suggesting that the helix is more stable in the dark state but ultimately unfolds at a similar rate to the wild-type protein. Based on these data, we hypothesize that the 1636 cm^−1^ transient can be assigned to the Q513 side-chain carbonyl itself, or at least that dynamics associated with this signal directly report on the hydrogen bond between N414/Q414 and Q513.

### ^15^N/^1^H-HSQC NMR Reveals the Jα helix is primed for unfolding in N414A AsLOV2

As the TRIR and FTIR are difference experiments that show changes in the protein due to light activation, we used ^15^N-HSQC NMR to characterize the Jα helix in wild-type, N414A, and N414Q AsLOV2 in both dark and light states (**Figure S7, S8, and S9**, respectively). Excitation of wild-type AsLOV2 with 488 nm light reveals the formation of cross peaks between resonances at 7.5 and 8.5 ppm, consistent with the data reported by Harper et al.^9^ In N414A AsLOV2, some of these cross peaks are already formed in the dark state (**Figure S8**), suggesting that N414A AsLOV2 is in a structurally primed state for photoactivation with some residues in active or partially active conformations. The 2D NMR spectrum of N414Q AsLOV2 resembles wild-type-AsLOV2 in that there is clear separation between peaks from 7.5 and 8.5 ppm that become more poorly resolved upon light activation (**Figure S9**).

The indole side chains of W490 and W557 have been previously shown to be sensitive to the structure of the Jα helix.^9^ W490 is located near the N-terminal end of the Jα helix while W557 is located on the Jα helix itself. Wild-type AsLOV2 shows a clear change in the chemical shifts of both Trp residues between the dark and light states (**Figure 5A**). The W490 indole side chain N-H has chemical shifts of 10.09 ppm (^1^H) and 127.75 ppm (^15^N) in the dark state and 10.12 ppm (^1^H) and 128.6 ppm (^15^N) in the light state, whereas W557 has chemical shifts of 10.26 ppm (^1^H) and 129.25 ppm (^15^N) in the dark state and 10.10 ppm (^1^H) and 129.00 ppm (^15^N) in the light state. In the dark state of N414A AsLOV2 there is a population of W557 that has chemical shifts similar to those observed in the light state while W490 has a similar chemical shift, albeit shifted to 128.00 ppm (^15^N), compared to wild-type AsLOV2 (**Figure 5B**). This suggests that the C-terminal end of the Jα helix is slightly unfolded due to mutation of N414 while the hinge-loop region connecting the Jα to the LOV core is intact. The disorder of the Jα helix in the N414A AsLOV2 dark state observed in these HSQC spectra is consistent with the smaller 1626 cm^−1^ bleach in the TRIR experiment suggesting that the helix is partially unfolded in the dark state.

**Figure 5:**
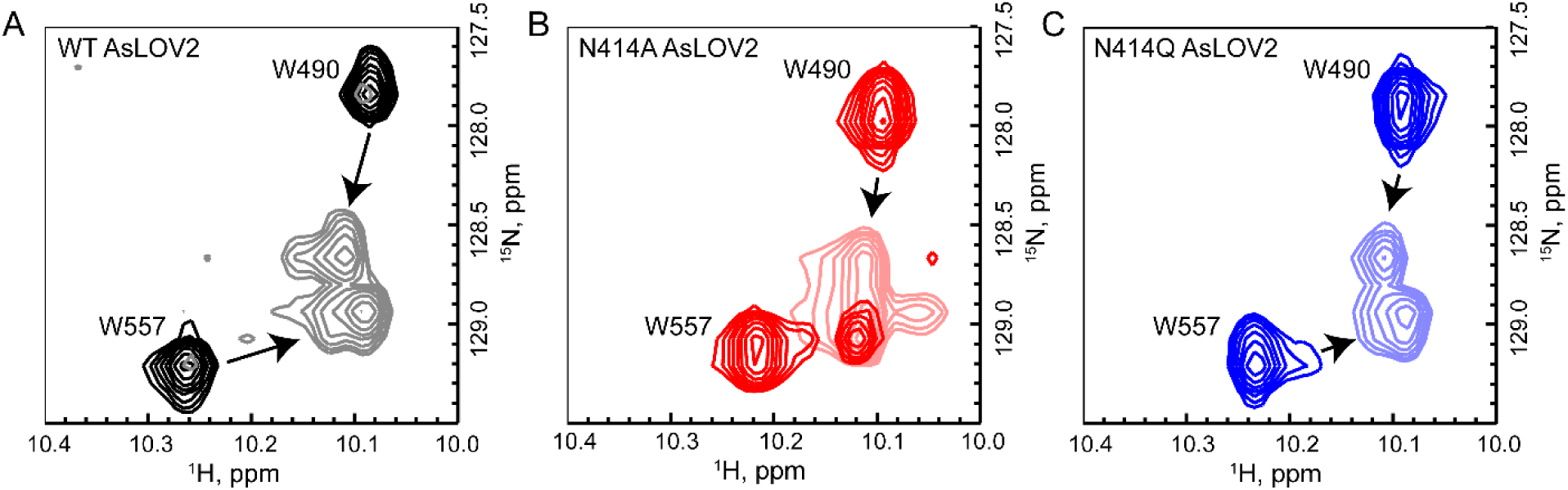
^15^N/^1^H-HSQC NMR reveals that the Jα helix is partially unfolded in N414A-AsLOV2. (A) Tryptophan indole side-chain chemical shifts of the wild-type protein show a clear shift and broadening from dark to light states (black to gray). (B) The same resonances in N414A AsLOV2 show that W557, which is located on the Jα helix, is partially in the lit state suggesting that the helix is partially unfolded (Dark red to light red). (C) N414Q AsLOV2 does not show partial unfolding, however the peak assigned to W557 is distorted compared to wild-type AsLOV2 (Dark blue to light blue).

### A cell-based assay shows loss of function associated with changes in Jα helix unfolding of the N414 mutants

To assess the functional consequences of mutating N414, we used the cell-based LOVTRAP system to measure the ability of AsLOV2 to dimerize to the plasma membrane localized Zdk2 protein and release from the plasma membrane upon light activation (**Figure 6A**).^34^

**Figure 6:**
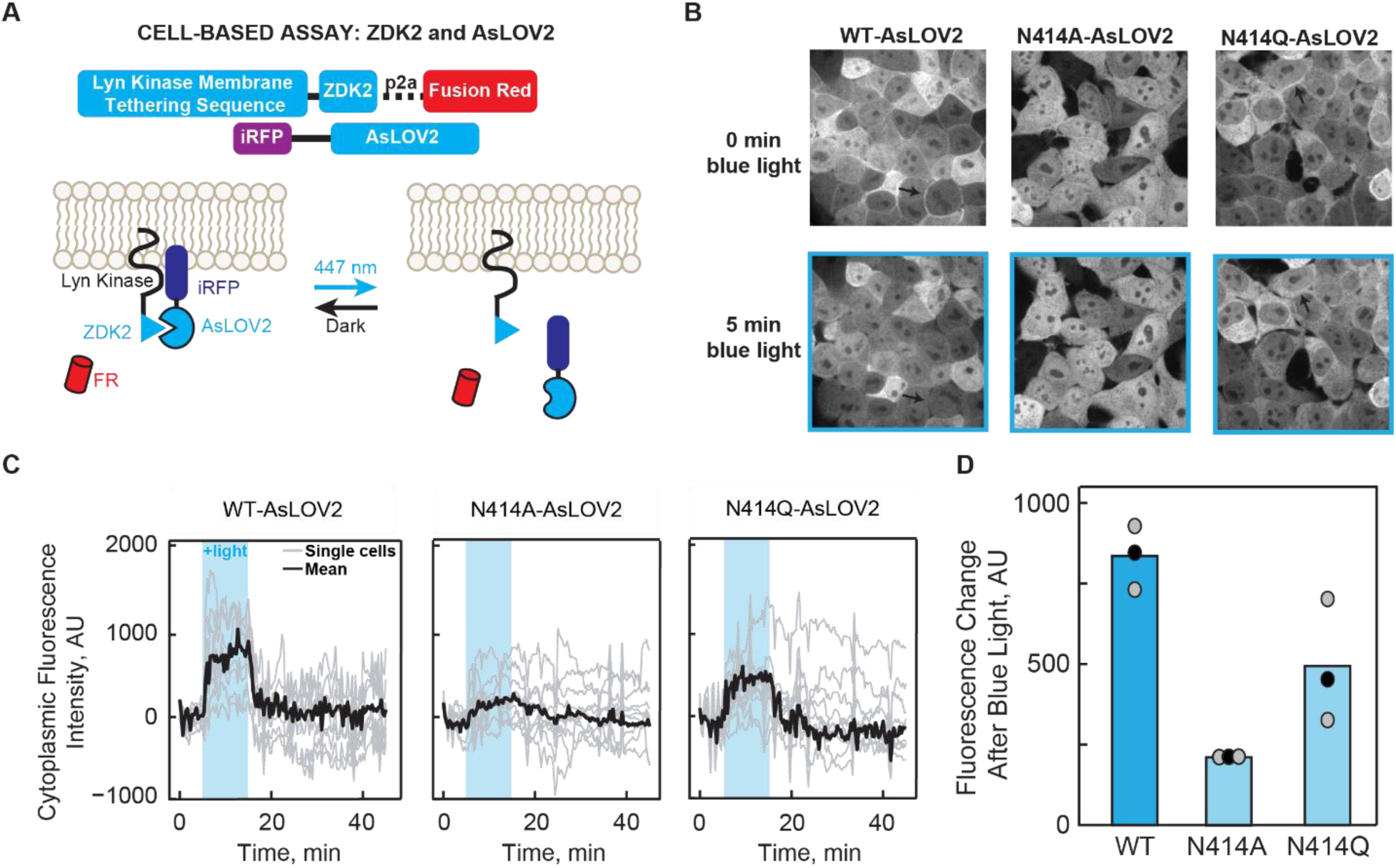
LOVTRAP assay shows altered Jα helix dynamics in N414A AsLOV2. (A) Schematic diagram of the LOVTRAP system. The Zdk2 domain binds to the Jα helix of AsLOV2 in the dark state and is released upon illumination with 447 nm light. The translocation of AsLOV2 is tracked using iRFP. (B) Representative images showing the localization of AsLOV2 to the plasma membrane in the dark state and diffusion into the cytoplasm after 5 min of blue light illumination. Localization on the plasma membrane is shown by black arrows. Fluorescence localization to the cell membrane can be observed in wild-type AsLOV2 and N414Q AsLOV2 but not N414A AsLOV2 in the dark state. Fluorescence localization to the cell membrane is not observed for the AsLOV2 variants after blue-light irradiation. (C) Quantification of the change in fluorescence in the cytoplasm over time shows that N414A AsLOV2 has reduced light-induced localization and ΔF compared to wild-type and N414Q AsLOV2. (D) Quantification of the total maximum change in fluorescence due to release of AsLOV2 from the membrane. Each point represents the average of 10 cells after 8 minutes of blue light exposure. The mean and standard deviations of each bar are shown as black and gray dots, respectively.

Localization of AsLOV2 to the plasma membrane is visualized by fusion of iRFP to the N-terminus of AsLOV2. For wild-type and N414Q AsLOV2, localization of the fluorescence signal is observed on the plasma membrane in the dark state and is released into the cytoplasm upon blue-light illumination (**Figure 6B**, black arrows denote plasma membrane) whereas the N414A variant shows minimal localization to the plasma membrane even in the dark state. This suggests that prior to illumination, N414A AsLOV2 exhibits light state activity. Quantification of the change in fluorescence intensity in the cytoplasm over time is shown in Figure 6C for all three AsLOV2 variants. The largest change in fluorescence intensity (ΔF) after illumination is observed in wild-type AsLOV2 with a mean ΔF of 835±57 units, while N414Q AsLOV2 shows a slight (1.7-fold) reduction in ΔF (493±110 units). These results are consistent with the NMR spectra in which the chemical shift of W557 in N414Q AsLOV2 shows more disorder compared to wild-type AsLOV2. N414A AsLOV2 shows a more dramatic 4-fold reduction in ΔF of 209±5 (**Figure 6D**) consistent with minimal membrane localization in the dark state. The reduced ΔF is indicative of reduced dynamic range in N414A AsLOV2, either a result of the partially unfolded helix in the dark state causing residual light state activity or a deficient unfolding of the Jα helix.

The decay of the fluorescence signal was fit to a single exponential function to determine the rate of dark state recovery of the LOVTRAP construct and was found to be 60 s ± 10 s and 205 s ± 28 s for the wild-type and N414Q AsLOV2 proteins, respectively (Supplemental Figure S10). These time constants correlate with the solution measurements of Zayner et al.^25^ in which rates of 80 s and 280 s were observed for wild-type and N414Q. N414A was not included in our analysis due to the small change in fluorescence between dark and light states (ΔF). The faster rate of recovery in the optogenetic experiment is likely due to the increased temperature from 22°C in the solution experiment to 37°C in the cell-based assay.

## Discussion

Photoactive proteins convert light energy into structural changes that drive and control biological processes by modulating the activity of downstream output partners.^35^ In the LOV domain proteins the light absorbing chromophore is the isoalloxazine ring of a non-covalently bound FMN cofactor. Excitation of the chromophore triggers a photocycle in which an early event is the formation of a covalent adduct between the FMN and a conserved Cys residue (C450) and protonation of FMN-N5.^3^ Cys-FMN adduct formation then results in slower time-scale structural changes that are transmitted through a C-terminal helix, the Jα-helix.^9^ In AsLOV2 the helix unfolds on the micro-millisecond timescale, and there have been numerous efforts to determine how the Cys-FMN adduct formation modulates the structure of the helix which is ∼13 Å away.^8^ In the current work we use a combination of MD, TRIR spectroscopy, NMR spectroscopy, site-directed mutagenesis and cell-based experiments to elucidate the pathway through which the chromophore and helix communicate.

X-ray structural studies have revealed that a conserved Gln, Q513, rotates during photoactivation. In the dark state, the side-chain NH_2_ and main chain C=O of Q513 are hydrogen bonded to the FMN C4=O and side chain of N414, respectively, whilst in the light state, the side chain carbonyl of Q513 now accepts a hydrogen bond from the protonated FMN-N5 (**Figure 1**). Rotation of Q513 is clearly a key event in light state formation since replacement of this residue with any other amino acid, even Asn, results in loss of photoactivity, and a series of studies have shown that this motion of Q513 is coupled to later events of the photocycle, beyond 20 µs.^30^ For example, whilst the bleach corresponding to Jα helix unfolding is not observed by steady state FTIR difference spectroscopy in the Q513L variant,^11^ studies by our group using TRIR spectroscopy have shown that the early steps of the photocycle (<10 µs) are not affected by mutation of Q513 to Ala. Previous MD simulations have provided additional insight into the role of Q513, suggesting the formation of a hydrogen bond between the Q513 side chain C=O and the N414 side chain N-H group. However, these simulations were limited to 1 µs and were not able to capture the unfolding of the Jα helix.^13^

In the present work we have extended the MD simulations to 7 µs which is sufficient to capture unfolding of the Jα helix and provide an atomic-level prediction of the events leading to light state formation. The simulation predicts a 62° rotation of the Q513 side chain leading to the formation of a transient hydrogen bond between the Q513 and N414 side chains 1.15 µs after light absorption, which is accompanied by a 3.1 Å movement of N482 out of the FMN binding pocket and a 31° rotation of N492, consistent with previous MD simulations.^13^ Unfolding of the Jα helix occurs over the lifetime of the transient Q513-N414 hydrogen bond, and the subsequent reorganization of key residues in and around the FMN binding site provide novel insights into the allostery of LOV domain activation in which N482 returns to its initial conformation whilst N492 remains rotated out of the flavin binding pocket and does not recover on the timescale of the simulation. While the H-bond network partially recovers by the end of the simulation, the Jα helix remains unfolded. Thus the 7 µs simulation provides insight into the complete photoactivation pathway by visualizing how the motion of the amino acids trigger larger secondary structural dynamics.

Recent structural studies of OdAureo1a bound to 5-deaza-FMN have also shown evidence of hydrogen bonding between Q513 and N414 (**Figure S11**), however the protein was found to be unable to dimerize as C5 in 5-deaza-FMN cannot be protonated and was therefore photoinactive.^36^ Using TRIR and NMR, we have shown experimentally that the N414 amide side chain is a key regulator of Jα helix unfolding through site-directed mutagenesis. In the N414A mutant, the TRIR showed a diminished bleach at 1626 cm^−1^ assigned to Jα helix unfolding (**Figure 3B,E**) and the time constant for the structural evolution from A390 to the final signaling state was accelerated ∼3-fold (Figure 4A). In contrast the N414Q mutant, which retains the amide side chain, is similar to wild-type AsLOV2. Since the TRIR experiment is a difference experiment and does not explicitly reveal the absolute dark and light state structures, we used NMR spectroscopy to determine the structure of the Jα helix in the dark and light states of N414A and N414Q AsLOV2 using the indole side chain N-H groups of W491 and W557 as probes.^10^ The HSQC NMR data showed that the W557 side chain in the N414A AsLOV2 dark state was populating the light state conformation and thus that the Jα helix is already partially unfolded in the dark state explaining the smaller change in the TRIR difference spectrum for this mutant (**Figure 5**).

In order to link the structural dynamics revealed by the MD simulations and spectroscopic studies with LOV domain function, we assessed the impact of the N414 mutations on the activity of an optogenetic LOV domain construct in the cell-based LOVTRAP assay. In the dark state, AsLOV2 is localized to the cell membrane due to interactions between the Zdk2 peptide/Lyn Kinase Transmembrane domain fusion and the Jα helix of AsLOV2 while in the light state, unfolding of the Jα helix causes dissociation of the complex and diffusion of AsLOV2 into the cytoplasm, which is visualized by iRFP fluorescence. While the wild-type and N414Q construct show localization to the cell membrane in the dark state and dissociation from the membrane upon illumination with blue light, the N414A mutant is minimally localized to the membrane in the dark state consistent with a reduction in the change in iRFP fluorescence upon illumination (**Figure 6**). These results directly implicate N414 as a key regulator of LOV domain function by coupling initial structural changes around the chromophore to Jα helix unfolding and photoactivation.

Taken together, our studies reveal that N414 plays two roles in the photoactivation dynamics of AsLOV2, by directly controlling the structure of the Jα helix in the dark state, and by coupling local structural changes around the FMN chromophore on light absorption with Jα helix unfolding. In the dark state, the side chain of N414 is hydrogen bonded to the backbone amide of Q513.^8^ This appears to stabilize the interaction between A’α and Jα, which was previously shown by Zayner and coworkers to be important for the unfolding of the Jα helix as observed by FTIR.^28^ The TRIR and NMR spectra show that N414 is responsible for stabilizing the Jα helix in the dark state and that this helix is partially unfolded in the N414A mutant which has a critical impact on LOV domain function since the N414A LOVTRAP mutant has lost the ability to interact with the Zdk2 peptide in a light dependent manner.

N414 also modulates the kinetics protein evolution that occurs between Cys-FMN adduct formation (A390) and Jα helix unfolding (Sig). The rise and decay pattern observed in the 1635 cm^−1^ band in the TRIR is consistent with the formation and breakage of structures involving amide carbonyl groups. Since these kinetics are not observed in N414A, we assign the 1635 cm^−1^ band to protein dynamics initiated by the transient H-bonding between Q513 and N414 and by extension N482 which is predicted to slide out of the FMN binding pocket and recover back to its initial H-bonding environment as Jα helix unfolds. Several different time constants have been reported for the rate at which Jα helix unfolds. Transient grating (TG) spectroscopy, which is a diffusion dependent signal, demonstrated that the helix unfolds in 2 steps in which the Jα helix undocks from the β-sheet in 90 µs and has fully unfolded in 1-2 ms.^37,38^ Here, we report a 313 µs time constant for Jα helix unfolding which is complete by 1 ms based on comparison of the TRIR and steady state FTIR difference spectra (**Figure 7, Supplementary Figure S4**) and is comparable to previously reported value of 240 µs from Konold et al.^29^ In contrast the Sig EAS of the N414A mutant evolves more rapidly with a 93 µs time constant and is consistent with Q513 rotating back to H-bond to FMN C4=O and N482 sliding back to H-bond to FMN C2=O faster than wild-type. Therefore, we propose a novel role of N414 in which the side chain modulate the longer µs kinetics of the LOV activation pathway.

**Figure 7:**
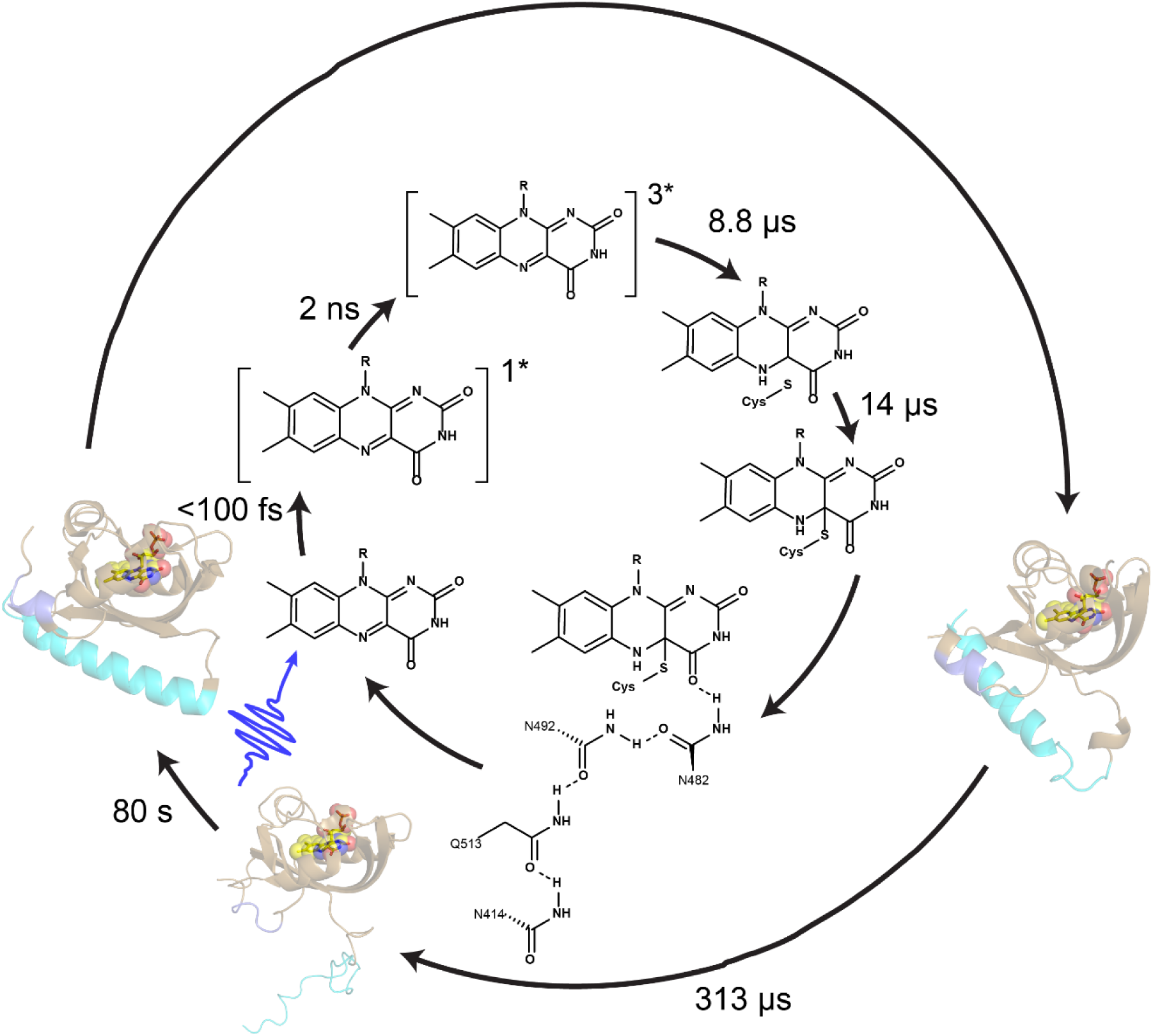
Phases of AsLOV2 activation with experimental time constants. Chromophore dynamics primarily occur on the ultrafast timescale while changes in the protein matrix occur much later with full Jα helix unfolding occurring with a time constant of 313 µs.

## Conclusion

In conclusion, we have used multiple approaches to shed new light on key events in the AsLOV2 photocycle immediately following Cys-FMN adduct formation, which occurs on the µs timescale and is associated with changes in the protein matrix at sites that are distant from the primary photochemistry. MD simulations reveal the formation of a transient hydrogen bond between Q513 and N414, two conserved residues in the LOV2 domain family, upon rotation of Q513. TRIR and NMR studies show that the structural dynamics are decoupled from FMN excitation in the N414A AsLOV2 mutant where the Jα helix is partially unfolded in the dark state, leading to residual dark state activity. The structural studies have been complemented with a cell-based assay which substantiate the critical role that N414 plays in transmitting information from the chromophore to the Jα helix. Together these results represent a high-resolution picture of the inner workings of a photosensory protein in which a glutamine lever induces microsecond structural dynamics via H-bond pathways that link the FMN chromophore to the Jα helix.

## Funding

This study was supported by the National Science Foundation (NSF) (MCB-1817837 to P.J.T., MCB-1750637 to J.B.F.) and the EPSRC (EP/N033647/1 to S.R.M.). J.N.I was supported by a National Institutes of Health Chemistry-Biology Interface Training Grant (T32GM092714). J.T.C. was supported by the IMSD-MERGE Program at Stony Brook University (5R25GM103962-04). J.E.T. was supported by NIH-DP2EB024247. A.A.G was supported by NIH-F32GM128304. A.L. is a Bolyai Janos Research Fellow and was supported by OTKA NN113090. J.B.F. would like to acknowledge the Research Corporation for Science Advancement for support from a Cottrell Scholar Award.

## Acknowledgements

The authors are grateful to STFC for access to the ULTRA laser facility. NMR data presented herein were collected at the City University of New York Advanced Science Research Center (CUNY ASRC) Biomolecular NMR Facility.

## Table of contents figure

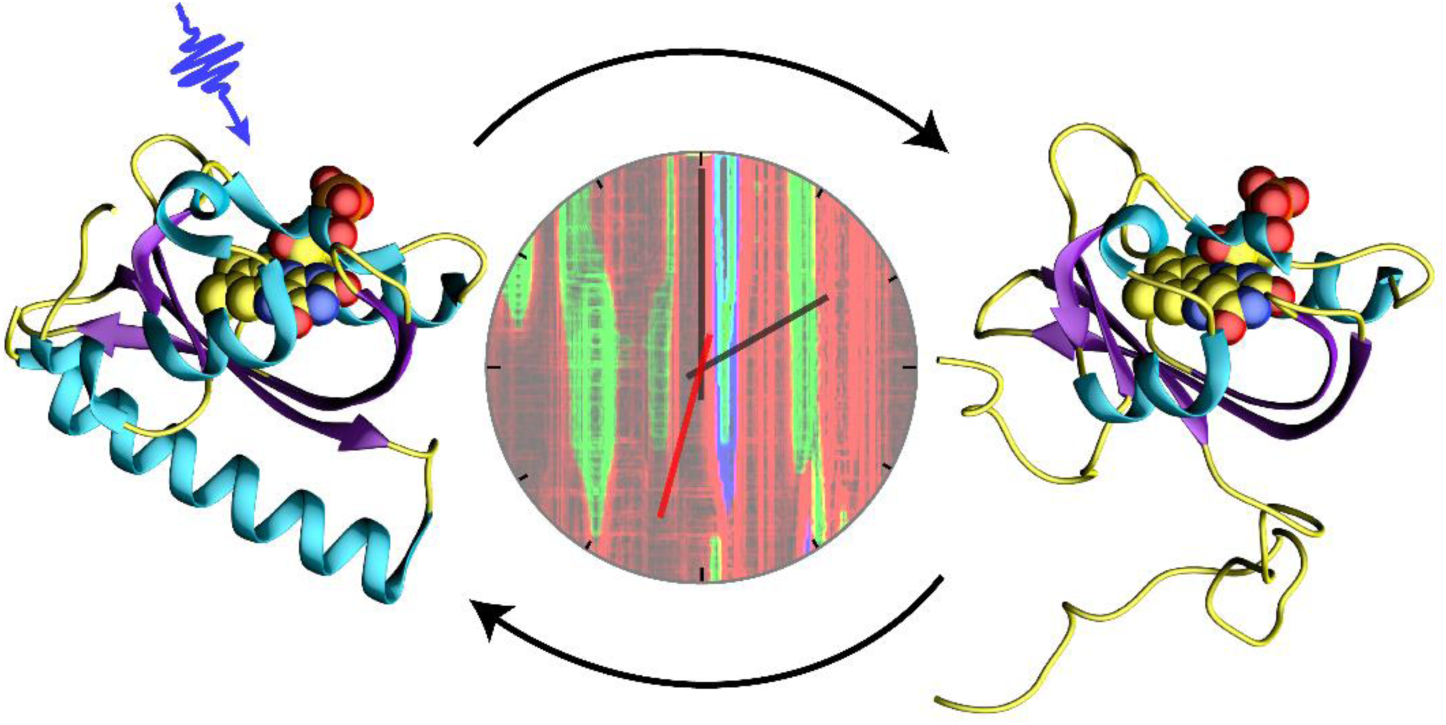

