## Supplemental Information for "Unraveling the Mechanism of a LOV Domain Optogenetic Sensor: A Glutamine Lever Induces Unfolding of the Jα Helix"

*[†]Department of Chemistry, Stony Brook University, New York, 11794, United States. [⊥]Department of Molecular Biology, Princeton University [‡]School of Chemistry, University of East Anglia, Norwich, NR4 7TJ, U.K. [§]Department of Biophysics, Medical School, University of Pecs, Szigeti út 12, 7624 Pecs, Hungary. [‖]Central Laser Facility, Research Complex at Harwell, Rutherford Appleton Laboratory, Didcot, OX11 0QX, U.K. [¥]Structural Biology Initiative, CUNY Advanced Science Research Center, 85 St. Nicholas Terrace, New York, NY 10031. [$]Ph.D. Program in Biochemistry, CUNY Graduate Center, New York, NY; [&]Ph.D. Programs in Biochemistry, Biology, and Chemistry, CUNY Graduate Center, New York, NY; [#]Department of Chemistry & Biochemistry, City College of New York, New York, NY; ; [±]Hormel Institute, University of Minnesota, Austin, MN, 55912.*

Keywords: LOV, structural dynamics, optogenetics, flavoprotein, ultrafast IR

**Table of Contents**


| | Page |
|---|---|
| Figure S1. Undocking of the Jα helix. | 3 |
| Figure S2: Comparison of light-state crystal structure and 6.8 us MD simulation. | 4 |
| Figure S3- Light-Dark FTIR Shows reduced helix unfolding in N414A-AsLOV2. | 5 |
| Figure S4: Evolution of difference spectrum on the 100s of μs timescale shows full structural changes in AsLOV2. | 6 |
| Figure S5: Kinetics of the C4a=C10a vibrational mode does not decay on the timescale of the experiment. | 7 |
| Figure S6: Selected traces showing quality of the global fitting. | 8 |
| Figure S7: HSQC NMR of WT AsLOV2 in both dark and light states. | 9 |
| Figure S8: HSQC NMR of N414A AsLOV2 in both dark and light states. | 10 |
| Figure S9: HSQC NMR of N414Q AsLOV2 in both dark and light states. | 11 |
| Figure S10: Fitting of the recovery traces from the LOVTRAP assay. | 12 |
| Figure S12: Comparison of 5d-FMN-Aureo with AsLOV2. | 13 |

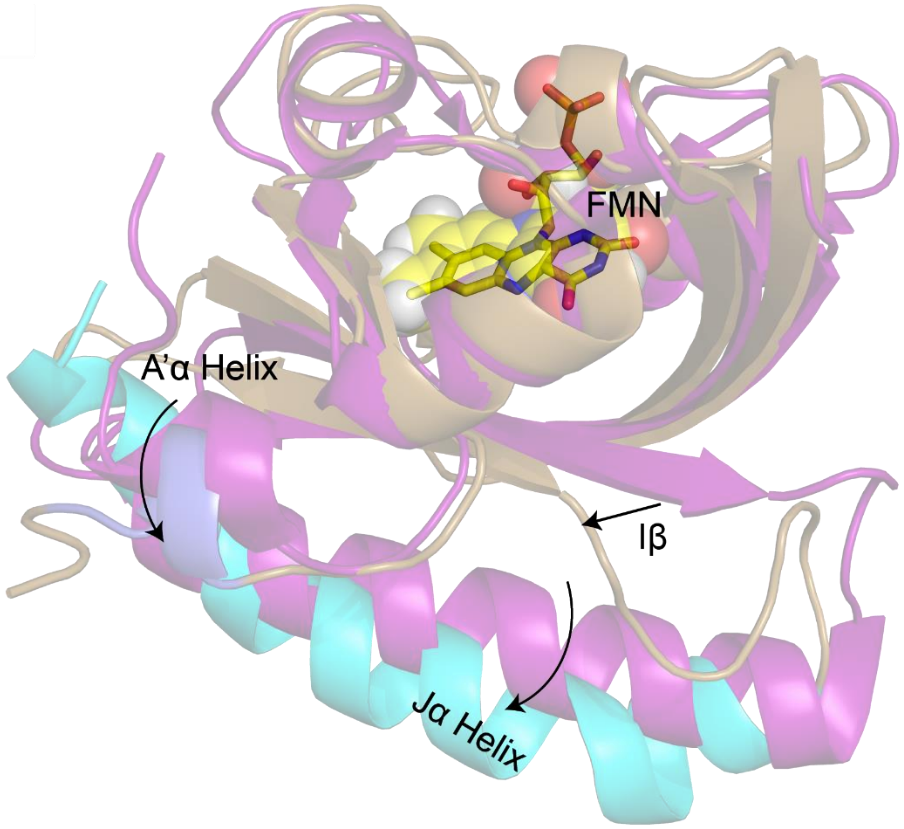


**Figure S1: Initial events of the Jα helix unfolding.** (A)The Jα helix is predicted to begin to undock ~0.5 μs after adduct formation. The dark state structure 2V1A is shown in magenta for reference. Iβ and the hinge loop region become more disordered allowing the Jα helix to begin to adopt an undocked conformation (cyan) and move away from the β-sheet (brown). The figure was made using the PyMOL Molecular Graphics System, Version 1.8, Schrödinger LLC.[19]

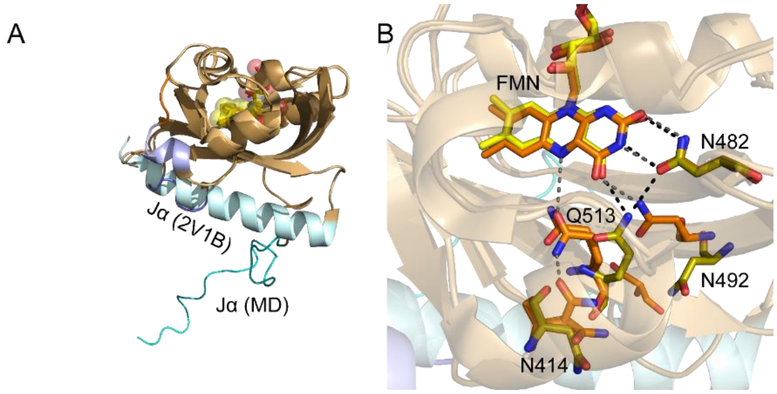


**Figure S2: Comparison of light-state X-ray structure and the structure formed after 6.8 μs of MD simulation.** (A) Helix unfolding is not observed in the light state structure (2V1B) due to crystal contacts but in the absence of these constraints becomes disordered following MD. (B) Increased mobility of the FMN H-bond network is predicted by MD simulations in which N492 is predicted to remain out of the pocket, 4 Å displaced from the position from the crystal structure. The figure was made using the PyMOL Molecular Graphics System, Version 1.8, Schrödinger LLC.[19]

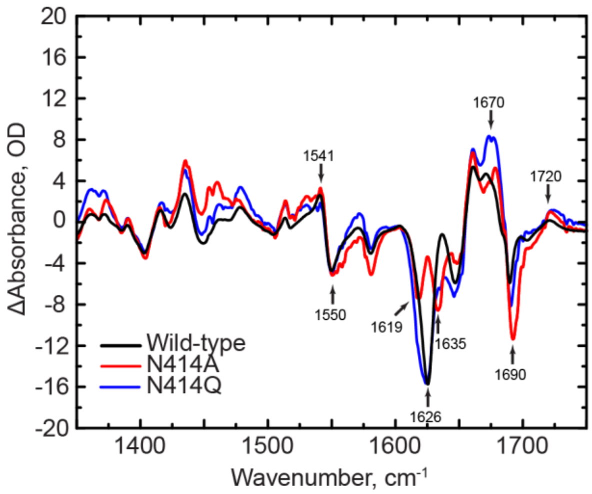


**Figure S3: Light-Dark FTIR Spectra of N414A-AsLOV2.** L-D FTIR difference spectra show full helix unfolding in WT-AsLOV2 and N414Q-AsLOV2 and reduced helix unfolding in N414A-AsLOV2 mutant. The marker band for the Jα helix is observed at 1626 $cm^{-1}$.

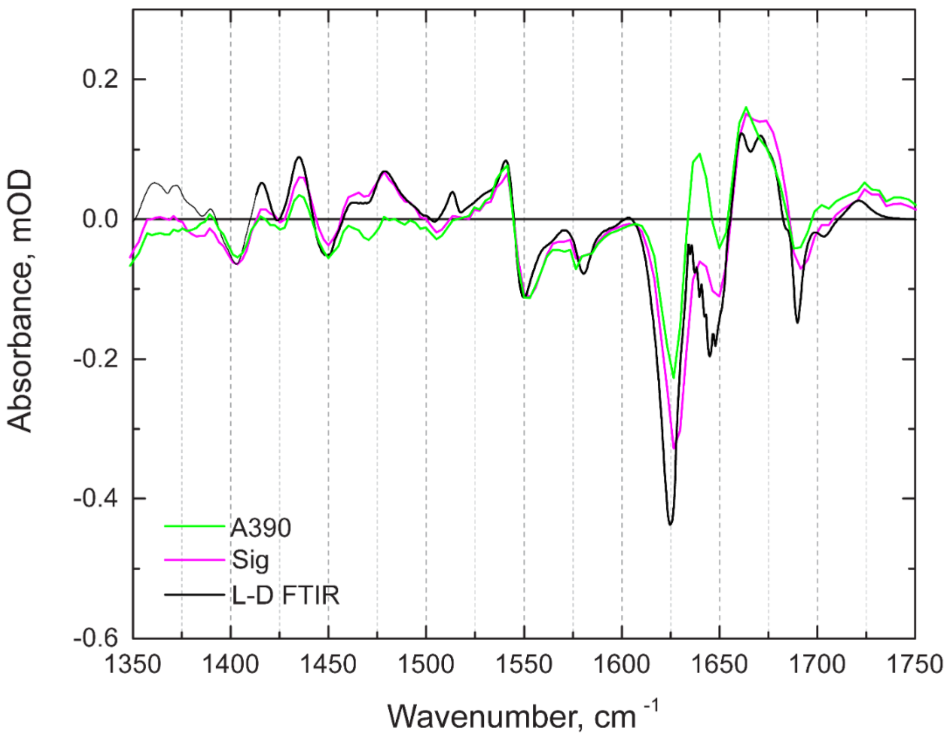


**Figure S4: Evolution of the FTIR difference spectrum on the 100s of μs timescale.** The full extent of the structural changes in wild-type AsLOV2 are shown by comparison of the light – dark FTIR difference spectrum (L-D FTIR, black) with EAS from a global fit of the TRMPS spectra corresponding to adduct formation (14 μs, A390 in green) and formation of the final signaling state (313 μs, Sig in purple). This data shows that full light state formation is captured in the TRMPS experiment.

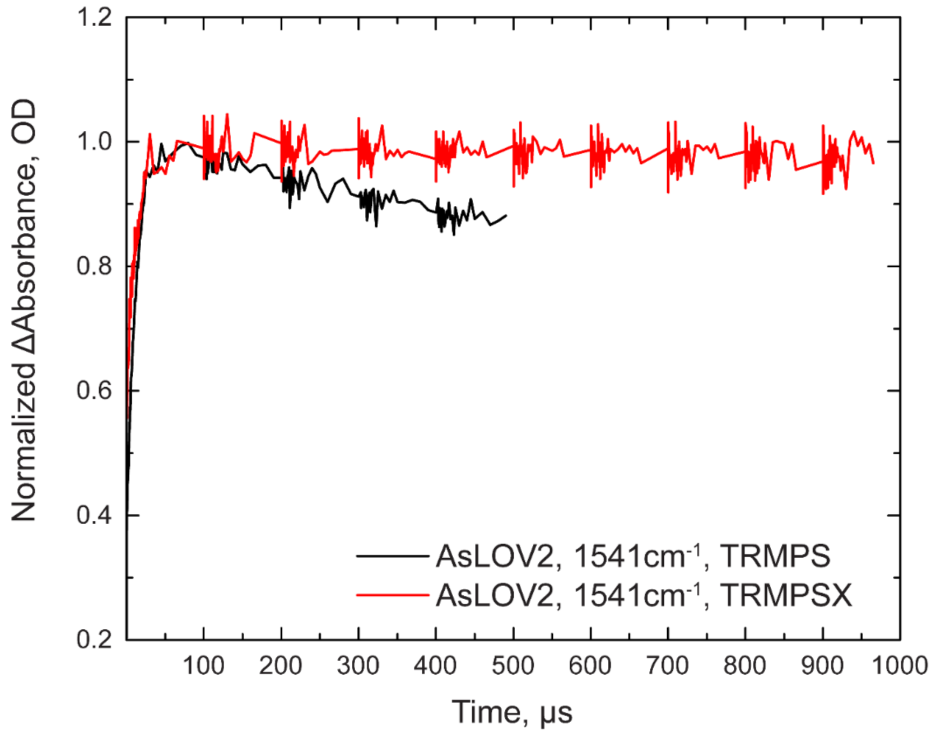


**Figure S5: Kinetics of the C4a=C10a vibrational mode from TRMPS and TRMPSX.** In the standard TRMPS experiment, a pump-off subtraction is taken at the final probe pulse and assumes that all signals are decayed by 1 ms. For AsLOV2 (and LOV in general), the light state is stable on the order of seconds such that signals from the light state are subtracted at later time delays. Employing the TRMPSX method and lowering the pump laser repetition rate to 500 Hz allows for a pump-off subtraction to be acquired at 2 ms, allowing for sufficient time for sample refresh prior to the pump-off pulse. The above traces compare the evolution of the 1541 $cm^{-1}$ transient assigned to the C4a=C10a vibrational mode from FMN acquired using TRMPS and TRMPSX. The signals acquired using TRMPSX do not decay on the ms timescale allowing for analysis of signals beyond 100 μs.

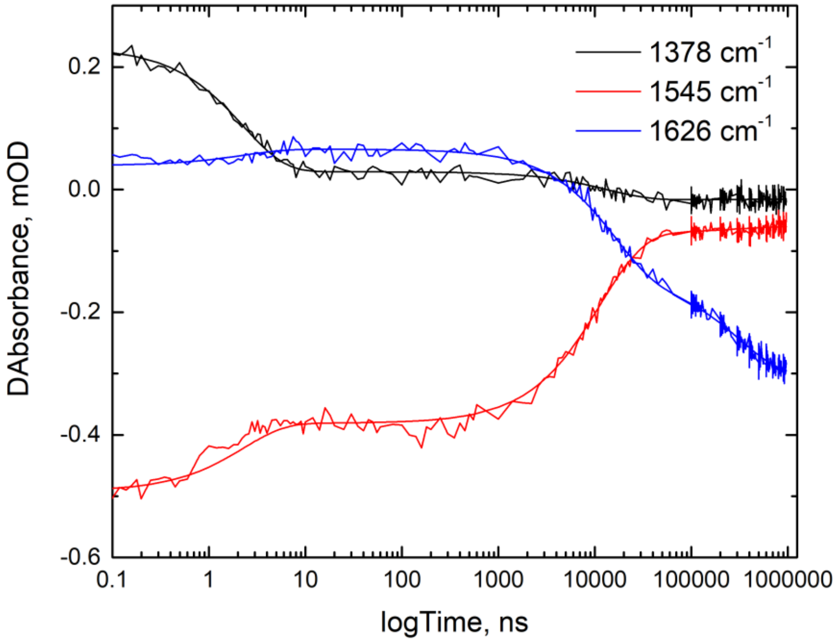


**Figure S6: Selected time traces showing quality of the global fitting.** Global fitting of the TRMPSX datasets to a sequential exponential model containing 5 components was used to extract time constants for the phases of evolution of AsLOV2 from dark to light state. Signals corresponding to excited state decay (1378 $cm^{-1}$), ground state recovery (1545 $cm^{-1}$), and the Jα helix (1626 $cm^{-1}$) are shown as representative kinetic traces.

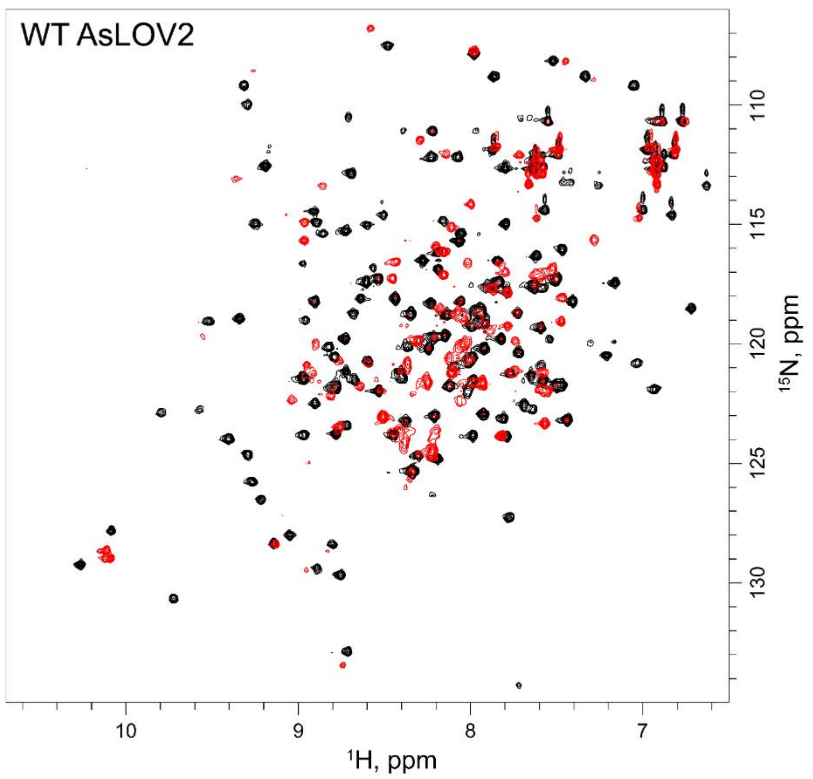


**Figure S7: $^{1}H$-$^{15}N$-HSQC NMR spectra of wild-type AsLOV2.** 800 MHz NMR spectra of $^{15}N$ labeled WT-AsLOV2 in the dark (black) and light states (red) are shown. The light state was generated by irradiation with 50 mW, 120ms pulse duration of 488 nm light prior to collecting each transient spectrum. Data were processed and analyzed using Bruker TopSpin.

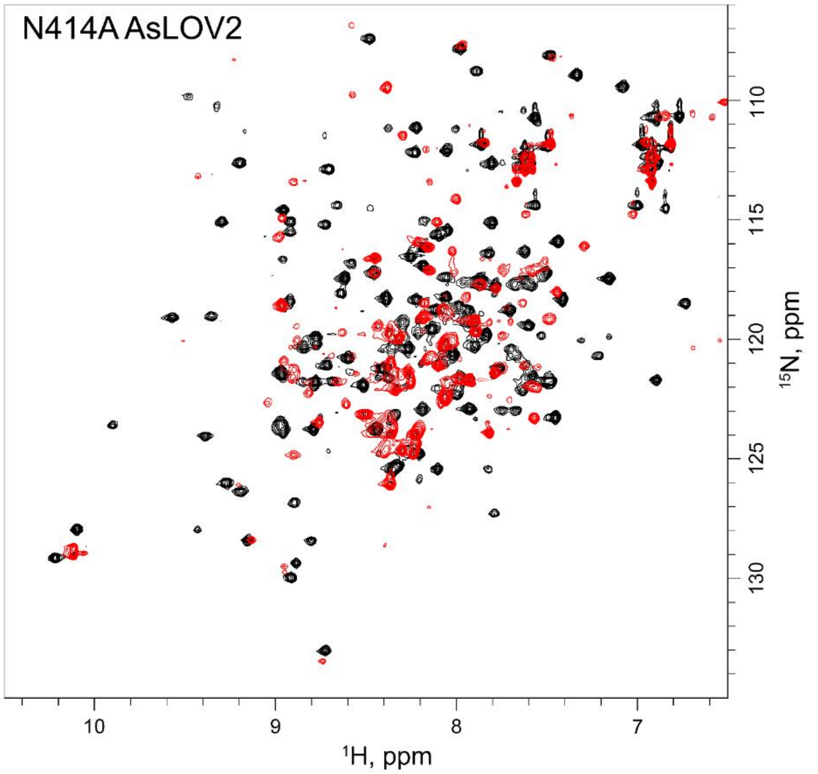


**Figure S8: $^{1}$H-$^{15}$N-HSQC NMR spectra of N414A AsLOV2.** 800 MHz NMR spectra of $^{15}$N labeled N414A-AsLOV2 in the dark (black) and light states (red) are shown. The light state was generated by irradiation with 50 mW, 120ms pulse duration of 488 nm light prior to collecting each transient spectrum. Data were processed and analyzed using Bruker TopSpin.

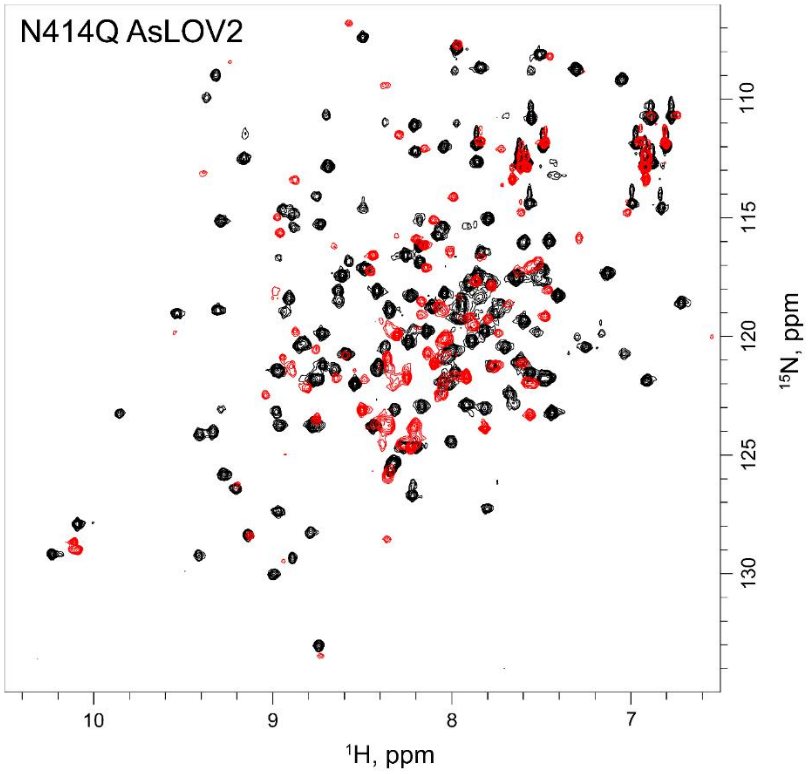


**Figure S9 : $^{1}$H-$^{15}$N-HSQC NMR spectra of N414Q-AsLOV2.** 800 MHz NMR spectra of $^{15}$N labeled N414Q-AsLOV2 in the dark (black) and light states (red) are shown. The light state was generated by irradiation with 50 mW, 120ms pulse duration of 488 nm light prior to collecting each transient spectrum. Data were processed and analyzed using Bruker TopSpin.

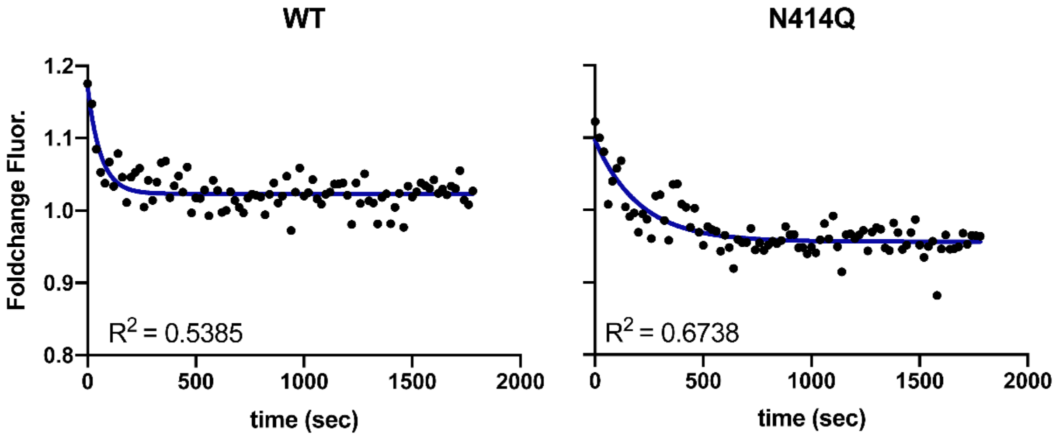


**Figure S10: Dark state recovery of AsLOV2 using the LOVTRAP assay.** The ΔF signal from after cessation of blue light irradiation from the time trace in **Figure 6** of the main text was fit to a single exponential decay to extract the recovery time of AsLOV2. The readout of ΔF in this assay is directly proportional to recovery (refolding) time of the Jα helix rather than recovery of oxidized FMN and is more reflective of protein dynamics. The time constants were determined to be 60 s ± 10 s and 250 s ± 28 s for the wild-type and N414Q-AsLOV2 proteins, respectively, which are consistent with recovery times measured in solution using UV-Vis spectroscopy.[20] N414A-AsLOV2 was not included in this analysis as the curve could not be adequately fit.

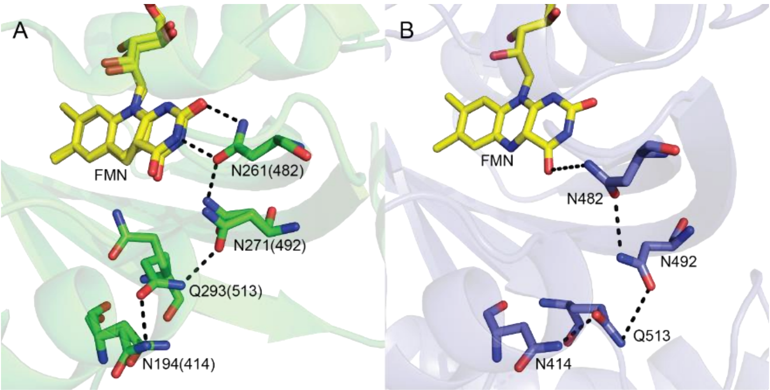


**Figure S11: Comparison of 5d-FMN-Aureo and AsLOV2.** (A) The crystal structure of the LOV domain of Aureo1a from *Ochromonas Danica (*OdAureo1a, PDB 6I24)[21] containing noncanonical FMN analogue, 5-deaza-FMN showed rotation of the conserved Q293(513) and formation of a stable H-bond between the Q293 C=O and the side chain N-H of N194. (B) The 1.15 μs snapshot from the MD simulation is shown for comparison. Q513 is predicted to be more mobile in the MD simulation and is 3.2 Å further from FMN in the 1.15 μs snapshot compared to the OdAureo1a structure. The figure was made using the Pymol Molecular Graphics System, Version 1.8, Schrödinger LLC.[19]
